# Joint inference of demography and mutation rates from polymorphism data and pedigrees

**DOI:** 10.1101/091090

**Authors:** Florence Parat, Sándor Miklós Szilágyi, Daniel Wegmann, Aurélien Tellier

**Affiliations:** Centre of Life and Food Sciences Weihenstephan, Technische Universität München, 85354 Freising, Germany; Department of Informatics, Faculty of Sciences and Letters, Petru Maior University, 540088 Tîrgu Mures, Romania; Faculty of Electrical Engineering and Informatics, Department of Control Engineering and Information Technology, Budapest University of Technology and Economics, H-1117 Budapest, Hungary; Department of Biology, University of Fribourg, Fribourg, Switzerland; Swiss Institute of Bioinformatics, Fribourg, Switzerland

**Author notes:** These authors contributed equally (Daniel Wegmann, Aurélien Tellier). Centre of Life and Food Sciences Weihenstephan, Technische Universität München, 85354 Freising, Germany.

**Keywords:** likelihood, domesticated species, sex-ratio bias

## Abstract

Inference of demography and mutation rates is of major interest but difficult because genetic data is only informative about the population mutation rate, the product of the effective population size times the mutation rate, and not about these quantities individually. Here we show that this limitation can be overcome by combining genetic data with pedigree information. To successfully use pedigree data, however, important aspects of real populations such as the presence of two sexes, unbalanced sex ratios and overlapping generations have to be taken into account. We present here an extension of the classic Wright-Fisher model accounting for these effects and show that the coalescent process under this model reduces to the classic Kingman coalescent with specific scaling parameters. We further derive the probability of a pedigree under that model and show how pedigree data can thus be used to infer demographic parameters. Finally, we present a computationally efficient inference approach combining pedigree information and genetic data summarized by the site frequency spectrum (SFS) that allows for the joint inference of the mutation rate, sex-specific population sizes and the fraction of overlapping generations. Using simulations we then show that these parameters can be accurately inferred from pedigrees spanning just a few generations, as are available for many species. We finally discuss future possible extensions of the model and inference framework necessary for applications to wild and domesticated species, namely the account for more complex demographies and the uncertainty in assigning pedigree individuals to specific generations.

---

Demographic processes shape the genetic variation and genetic structure of populations and species, but also affect the efficacy of selection. There is thus considerable interest in inferring past demographic events, not least to serve as a null model for the identification of markers under selection. To this end, several methods have been proposed to estimate past demography from genetic data, and many of which are based on coalescent theory using either likelihood methods (e.g., Hey and Nielsen 2007; Hey 2011; Excoffier *et al*. 2013) or simulations embedded in Approximate Bayesian Computation (e.g., Beaumont *et al*. 2002; Wegmann *et al*. 2009). In these approaches, the coalescent model is used to link the genetic information to demographic parameters by integrating over the unobserved genetic relationships between individuals in a population in a computationally efficient way.

While the standard coalescent approach assumes no prior information on the true (parent-offspring) relationship between genetic lineages, knowledge on this would bring complementary information and hence increase estimation accuracy. Indeed, several methods have been proposed to use pedigree information to infer demographic processes using the increase of inbreeding over time under a given reproduction model (Falconer and Mackay 1996; Gutiérrez *et al*. 2008). Additionally, a method that allows the simulation of pedigrees under a given demographic and reproductive model and draws genealogies inside these pedigrees was developed (Gasbarra *et al*. 2005), and could be used, in an ABC framework, for inference. However, there is currently no general inference framework for such data.

Here we develop a maximum-likelihood method to infer demographic parameters from both pedigree information and genetic data summarized by the site frequency spectrum (SFS) of a sample. We postulate that the pedigree of a sample contains information about the demography of the population at least partially complementary to the information contained in the genetic data and a method exploiting the full information should therefore improve the inference of demographic parameters.

An additional advantage of our framework is its ability to infer effective population sizes (*N*_*e*_) and mutation rates (*μ*) jointly. Under the standard coalescent framework, both parameters are simply scaling the coalescent tree and hence only their product can be estimated, usually in the form *θ* = 4*N*_*e*_*μ*. Wakeley and Takahashi (2003) showed that a joint estimation becomes feasible if the sample size n exceeds *N*_*e*_ since the rate of coalescence in the first few generation is a function of the ratio *n*/*N*_*e*_ and hence contains information about *N*_*e*_ regardless of *μ*. This was later used to infer gene specific mutation rates in humans from deep sequencing data Nelson *et al*. (2012); Schaibley *et al*. (2013). As we show here, pedigree information also contains information about *N*_*e*_ independent of *μ*, enabling the joint inference of both demography and mutation rate even in case where *n* ≪ *N*_*e*_.

Pedigree information is available for many populations or species, in particular for managed populations under conservation management or domesticated animals under active breeding (e.g., Clutton-Brock *et al*. 1982; Ellegren 1999; Cunningham *et al*. 2001; Mc Parland *et al*. 2007). Yet many of these species have important life history traits that are not reflected in the standard Wright-Fisher model, including overlapping generations and two sexes. Additionally, in most domesticated species, fewer males are reproducing than females but males can reproduce over a longer time period, spanning several generations. While such life history traits have an impact on the response of the population to selection and are therefore accounted for in breeding programs Hill (1974), changes in allele frequencies due to drift remain well described by scaling the models with an appropriate effective population size (*N*_*e*_) Wright (1931); Engen *et al*. (2007). As a consequence, demographic inference in domesticated species using coalescent theory does usually not model overlapping generation or sex biased population sizes.

However, simple scaling does not extend to models incorporating pedigree information. To address this, we present here a Wright-Fisher-based diploid two-sex model with overlapping generation well describing pedigrees observed from domesticated breeding programs. We then show how this model results in a simple scaling of the standard coalescent in the absence of pedigree information and derive some analytical and numerical solutions to obtain estimates of the model parameters using the information contained in the SFS and pedigree data jointly. This also allows for the estimation of important life history characteristics such as the degree of overlapping generations and sex specific population sizes from such data.

## The Model

To account for life history traits very common in animal populations, we extend the classic Wright-Fisher model to a diploid species with two sexes and overlapping generations. Our model, which is schematically depicted in Figure 1, assumes discrete generations consisting of *N*_*f*_ female and *N*_*m*_ male individuals. We further assume random mating in that in each generation, and going backward in time, each individual picks a random female and male individual of the previous generation as mother and father, respectively.

**Figure 1.**
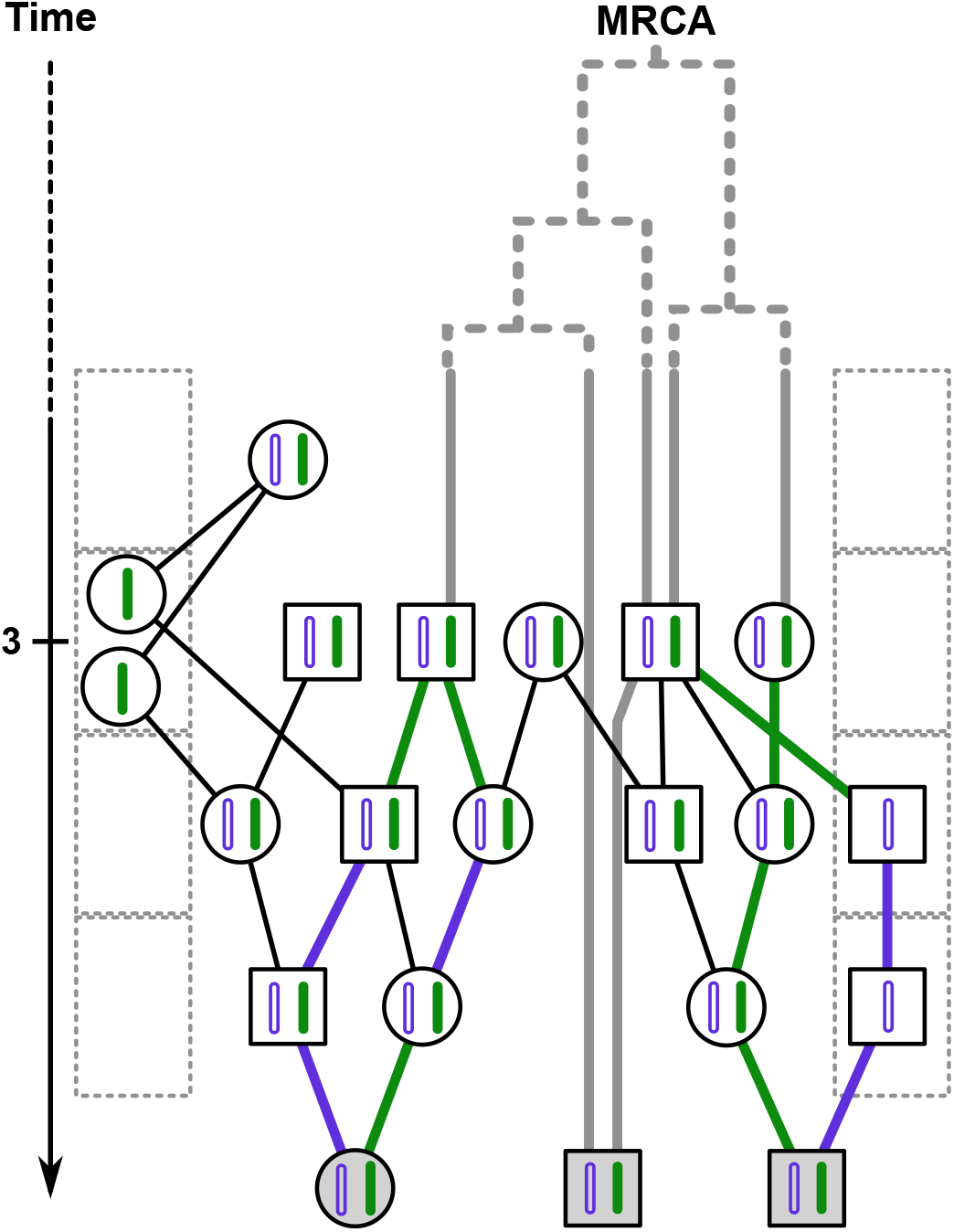
Graphical representation of the proposed two-sex model with overlapping generations. Shown are female (circles) and males (squares) individuals along with their parent-offspring relationships (black edges). The doted boxes on each side represent the female and male gamete storages populated with gametes of individuals from the pedigree. or the individuals of generation 0 (grayed) genetic data is available and the thick lines represent the genealogy of these individual. Dotted lines indicate the part of the genealogy before *g*_Max_ for which the continuous time approximation is used. The tick mark on the time scale represent the depth of this pedigree, *d* = 3.

We model overlapping generations by allowing individuals to pick their parents not from the directly preceding generation but from an earlier one with probabilities *b*_*f*_ and *b*_*m*_ for female and male parents, respectively. In this case, however, the choice of the actual distant parent is delayed and the lineage is just stored. In biological terms, these stored lineages thus represent gametes of a defined sex from previous generations, and we will refer to this compartment as “gamete storage” in the following. At the beginning of a generation, the so stored gametes then pick a parent in the current generation with probabilities 1 − *b*_*f*_ and 1 − *b*_*m*_, and otherwise remain in storage, which implies that the number of generations between parents of offspring are exponentially distributed.

For simplicity, we will considered here only the case of constant population sizes *N*_*f*_, *N*_*m*_ and probabilities to jump a generation *b*_*m*_, *b*_*f*_, but we note that relaxing this assumption is straight forward under the inference framework introduced below.

### Derivation of the coalescent

Two-sex models were previously shown (Möhle 1998) to be approximated accurately by the time-changed Kingman coalescent (1982). Similarly, Blath *et al*. (2013) recently showed that models with overlapping generations also result in a simple scaling of the classic coalescent if the average time lineages spend in storage is relatively small compared to the waiting time between coalescent events. Here we derive the appropriate scaling for the model introduced above.

We begin with the rate of coalescent and note that only the lineages that are in individuals of the real populations can coalesce, whereas lineages currently stored for later use have first to reenter the real populations. To obtain the fraction of lineages that can coalesce with each other, we obtain their relative distribution at equilibrium in the following four possible compartments: the real female (*R*_*f*_) and male (*R*_*m*_) populations as well as the female (*B*_*f*_) and male (*B*_*m*_) gamete storages (Supplementary Figure S1) using the following system of difference equations:

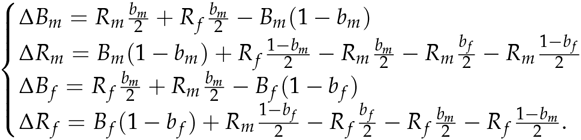

The global rate of coalescence Pr(Coal) is then given by sum of the per compartment rates weighted by the fraction of lineages residing in them. Since the coalescent rates are zero in the gamete storages and 1/2*N*_*f*_ and 1/2*N*_*m*_ in *R*_*f*_ and *R*_*m*_, respectively, we have

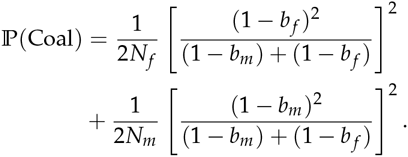

In accordance with previous results for two-sex (Möhle 1998) or seed bank (Kaj *et al*. 2001) models, the obtained rate reduces to

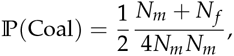

if *b*_*f*_ = *b*_*m*_ = 0, and to

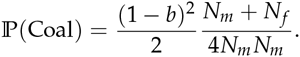

if *b*_*f*_ = *b*_*m*_ = *b*.

Following Kingman's approach, the distribution of time of coalescent under our model is 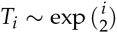 with time scaled in 2*N*_*e*_ with

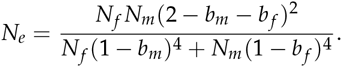

We next derive the rate of novel mutations in the presence of overlapping generations. Importantly, the number of germ-line mutations may not scale linearly with time, as, especially in females, most of these mutations occur during early development (Crow 2000). We model this effect using two mutation rates: *μ* for the part of the branches connecting real individuals and *μ* = εμ* for the time spent in the gamete storage. From the compartment model introduced above, we obtain the average fraction of time *t*_*b*_ that linages spend in one of the gamete storages as

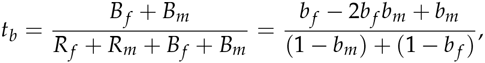

which results in an average effective mutation rate 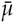 per generation:

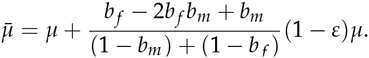

## Inference

We introduce here a Maximum Likelihood (MLE) method to infer jointly the demographic *θ*_*d*_ = {*N*_*f*_, *N*_*m*_, *b*_*f*_, *b*_*m*_} and mutational *θ*_*m*_ = {*μ, ε*} parameters of the model introduced above. This estimation is based on genetic data summarized by the site frequency spectrum (SFS) and available pedigree information in terms of child-parent relationships (filiation) that form one or several connected networks spanning two or more generations (𝒫). The relevant likelihood function can be decomposed as

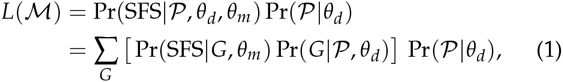

where the sum runs over the unknown genealogies *G* representing the genetic relationships between all sampled individuals up to the the most recent common ancestor (MRCA). While the pedigree and the genealogies share similar features, they should not be confused.

In the following sections, we will first derive each term of the likelihood function individually, and then give a detailed description of an inference framework under this model.

### The Pedigree

Let 𝒫_*g*_ be the way in which the individuals of generation *g* − 1 in the pedigree are assigned to their parents in generation *g*. Note that generations as well as the choice of the mother and the father are independent, and hence that

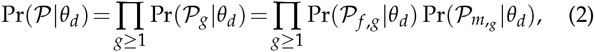

where 𝒫_*f,g*_ and 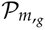 represent the assignment of individuals to their mothers and fathers, respectively.

The pedigree spans between the generation of the most recent individual *g* = 0 and the generation of the last known parent that we call *g*_Max_. To derive the probability of the pedigree, we consider all individuals at generation 0 in the pedigree as numbered (i.e. identifiable). These individuals then choose their parents from the previous generation, but are constrained in their choices by the pedigree. The so chosen parents become identifiable themselves and in turn choose the parents from the previous generation according to the pedigree information of that generation. This process continues until the top of the pedigree is reached.

Here we will derive Pr(𝒫_*f,g*_|*θ*_*d*_) for this process for the individuals of generation *g* − 1, of which exactly *B*_*f,g*−1_ will enter the gamete storage with probability *b*_*f*_ as their mother is from a distant generation. Among the 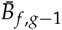 that choose a mother from generation *g*, the first individual of each of the *M*_*g*_ groups of siblings chooses a distinct mother from the population, which they do in turn with probabilities 1, 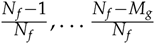. The *M*_*g*_ so chosen mothers, which have become identifiable themselves, are chosen by their remaining offspring with probability 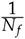 each. The resulting probability of this process is

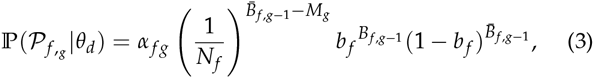

where we used the notation

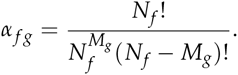

The same holds true analogously for ℙ(𝒫_*m,g*_|*θ*_*d*_) by replacing the subscript *f* by *m* and using *F*_*g*_, the number of fathers in generation *g*, instead of *M*_*g*_.

A maximum likelihood estimate of *N*_*f*_, *N*_*m*_, *b*_*f*_ and *b*_*m*_ is easily obtained by taking the first derivative of the logarithm eq. 2. For *b*_*f*_, thies yields

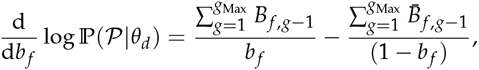

which admits the maximum likelihood estimate

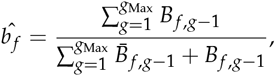

and analogously for *b*_*m*_. For *N*_*f*_, the first derivative is

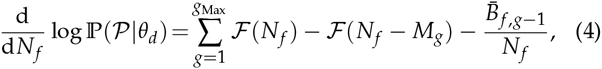

where 𝓕 is the digamma function defined as the logarithmic derivative of the factorial function. As the population size is a finite natural number, the maximum of this probability, if finite, can be easily found numerically using dichotomy.

### Genetic data

Coalescence is the merging of two or more genetic lineages. In a diploid population, an offspring may inherit one of two possible chromosomes of each parent. There are thus 2^*l*^ ways in which *l* offspring lineages can be assigned to the two chromosomes of a single parent (Figure 1). Enumerating all possible genealogies constraint by even a small but fully resolved pedigree, as done for two lineages in Wakeley *et al*. (2012), is computationally already very challenging for large sample sizes, and easily becomes prohibitive if the pedigree is only partially known. We thus chose to turn to simulations to evaluate the sum in eq. 1, as is commonly done in the absence of pedigree information (e.g. Excoffier *et al*. 2013; Nielsen 2000; Nelson *et al*. 2012):

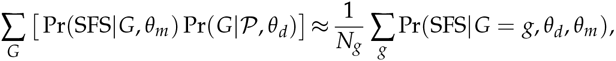

where the genealogies *g* ~ Pr(*G*|𝒫, *θ*_*d*_, *θ*_*m*_) are simulated under model parameters 𝓜 and constrained by the pedigree 𝒫.

Simulating genealogies inside a pedigree is straight forward and only requires binary choices when following lineages backward in time through the pedigree. In case of only partial pedigree information, lineages reaching parents of which only one or none of the parents are known choose their unknown parents randomly from the whole population, or enter the gamete storage (Figure 1). Since it is required to keep track of all these choices, the simulations become rather time consuming in case of limited pedigree information. At a certain generation in the past we term *g*_Max_, the pedigree does not contain any information about ancestors anymore and the genealogy is then only constraint by the parameters of the model. As *g*_Max_ is often reached long before the MRCA, we make use of the appropriately scaled coalescent approximation introduced above to simulate the genealogies from *g*_Max_ backwards to the MRCA (Figure 1).

To calculate Pr(SFS|*G* = *g, θ*_*m*_), the probability of the genetic data summarized by the SFS given a genealogy *g*, we use the classic infinite site mutation model with Poisson distributed mutations at rate 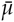 per site. Under this model, the probability that a mutation results in a derived sample allele frequency of *i* is given by the summed length *L*_*i*_ of all branches with *i* leaves and the probability of the SFS is thus given by a multinomial distribution

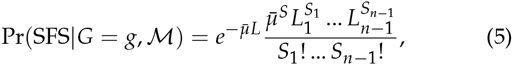

where *S*_*i*_ is the number of segregating sites being shared by *i* chromosomes in the sample of size *n* and L the total length of the genealogy g (Fu 1998). We note that the branch lengths *L*_*i*_ are measured in mutational generations, and hence all generations spent in the gamete storage only add *ε* to the total length.

The maximum likelihood estimate of 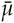 can be obtained analytically by differentiating the logarithm of eq. 5, which yields the estimator

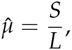

where L is the total length of the genealogy in mutational generations. In the absence of pedigree information, for instance for the part of the genealogy simulated under the coalescent approximation, the total length of the genealogy is only available measured in generations. In this case, the ML estimate becomes

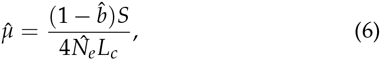

where *L*_*c*_ is the total length of the genealogy in coalescent time (i.e., in *θ* generations) and 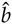 and 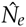 are the ML estimates of *b* and *N*_*e*_, respectively.

### Inference algorithm

An exact analytical or numerical solution for the joint maximum likelihood of all parameters is not available. We will therefore combine some of our analytical derivations with numerical evaluations to perform an MCMC-MLE inference as follows:

1. We sample vectors of demographic parameters 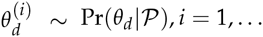, I from their joint posterior distribution given the pedigree using an MCMC framework.
2. For each sampled vector of parameters 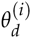, we simulate *G* = 100 genealogies constraint by the pedigree 𝒫.
3. For each 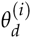 we then compute the MLE estimate 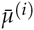 according to eq. 6 using the sampled genealogies.
4. Finally, we compute the joint likelihood of all model parameters for each pair of 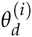 and 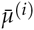 according to eq. 1, again using the simulated genealogies.

Our inference scheme is thus closely related to a grid search on the model parameters where we make use of the pedigree information to conduct the simulation based likelihood evaluation at promising locations only. The proposed combination of MCMC sampling and MLE is possible because the population size is constant and the maximum likelihood of 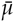 does only depend on the total length of the genealogies and not on their topology. As shown in the results, our MCMC-MLE method is an efficient compromise between speed and accuracy. Further advantages and limitations are discussed in the last section of the article.

The MCMC sampling in step 1 is implemented using a standard Metropolis algorithm (Metropolis *et al*. 1953) in which a single parameter is updated per iteration using a Gaussian proposal kernel mirrored at prior limits. We use uniform priors on all parameters except the population sizes for which we use loguniform priors and propose updates on the logarithmic scale during the MCMC to account for their prior easily spanning several orders of magnitude.

For all the runs presented here, we run the MCMC for 4.5 × 10^6^ steps thinned out to keep only every 500th parameter combination, of which the first 200 were discarded as a burn-in (resulting in 8800 sampled parameter vectors *θ*_*d*_). We use a normal distribution for the kernel of all three estimated demographic parameters (*N*_*f*_, *N*_*m*_ and *b*). The MCMC parameters relative to each of these demographic parameters can be found in the supplementary Table S1. The implementation of the method in C++ is available upon request.

## Simulations

To test the performances of our inference method, we used a custom R script to simulate pseudo observed datasets (PODs) consisting of a pedigree and a corresponding SFS for a sample of 50 individuals, unless specified differently. The pedigree includes all ancestors of the sampled individuals until the predefined depth *d* as well as the parents of all lineages in gamete storage at generation *d* (Figure 1). Thus, the generation of the oldest individual contained in the pedigree g_Max_ is such that *g*_Max_ ≥ *d*. We set *b*_*f*_ = *b*_*m*_ = *b* for all simulations and generated an SFS by simulating 2000 loci of 10 kb each with *μ* = 5 × 10^−9^ and *ε* = 0.

We first used simulations to assess the benefit of having pedigree data as a function of the pedigree depth across 10 independently generated PODs with demographic parameters realistic for domesticated breeds (*N*_*f*_ = 5000, *N*_*m*_ = 500 and *b* = 0.2). As shown in Figure 2, our method is capable of accurately disentangling the effects of the mutation rate and population sizes on genetic diversity already if limited pedigree information is available. Indeed, reliable estimates are obtained for all parameters including sex-specific population sizes, the frequency of overlapping generations as well as the mutation rate if a pedigree of depth four or more is used (Figure 2).

**Figure 2.**
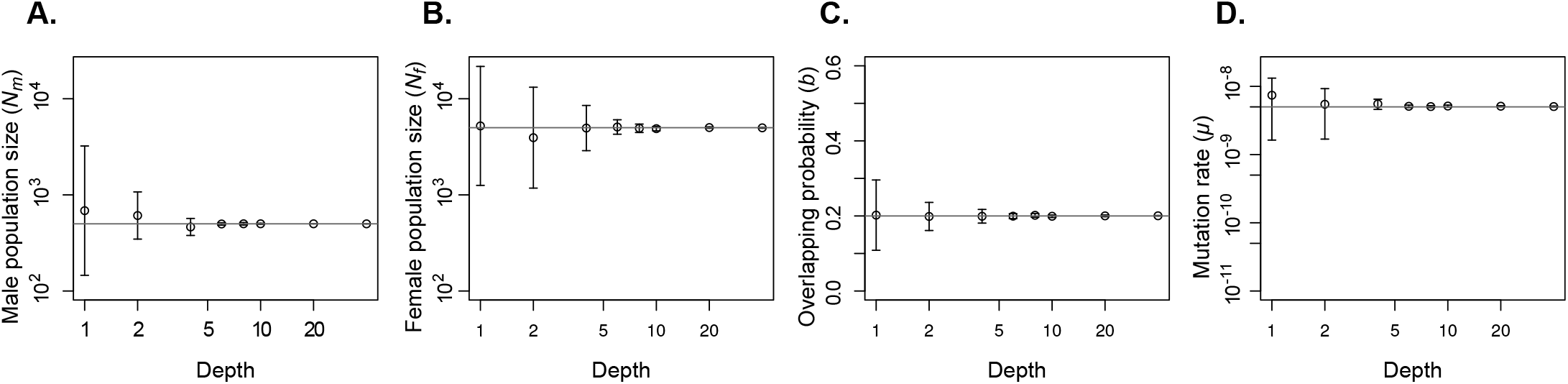
Power of parameter inference as a function of pedigree depth. Shown are the mean and standard deviation over 10 simulated datasets. The true parameter values used for all simulations (A) *N*_*m*_ = 500, (B) *N*_*f*_ = 500, (C) *b* = 0.2, and (D) *μ* = 5 × 10^−9^ are indicated by gray horizontal lines.

Interestingly, the rate of overlapping generations *b* is estimated well across the whole parameter range in the presence of sufficient pedigree data (Figure 3), but smaller population size seem to be consistently estimated more accurately than lager sizes. This is visible as a reduced accuracy in the inference of the female compared to male size in Figure 2, but also occurs if the population sizes of both sexes are equal (Figure 3 and 4). We explain this as follows. The information about the population size of a pedigree is mostly contained in individuals sharing parents (i.e., half or full siblings). If the population is large but the number of individuals in the pedigree relatively small, few to no siblings are observed and the power to estimate the population size decreases. Indeed, when there are no siblings in the pedigree, the likelihood of the population size increases monotonously but reaches a kind of plateau before the true value is reached (e.q. 4). This leads to an overestimation of the population size in the absence of genetic information and the inability to disentangle *N*_*e*_ from *μ* if such data is available. As an example, consider the posterior distributions shown in Supplementary Figure S2 for the case of *N*_*f*_ = 5,000 and a pedigree depth of one.

**Figure 3.**
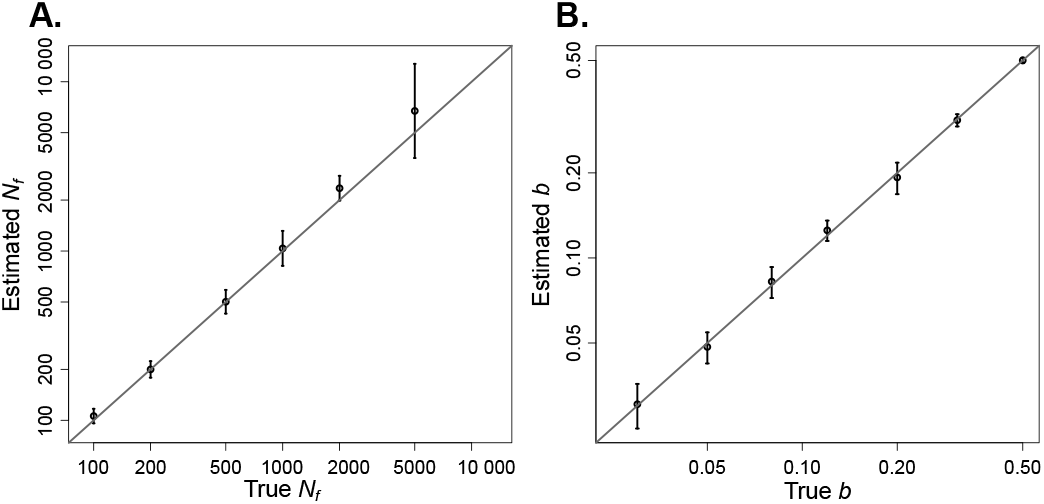
Estimated (A) N_e_ in function of simulated N_e_ and (B) b in function of simulated b, in log-log scale. The vertical bares represent the standard deviation over 10 datasets. The gray diagonal marks the identity line.

**Figure 4.**
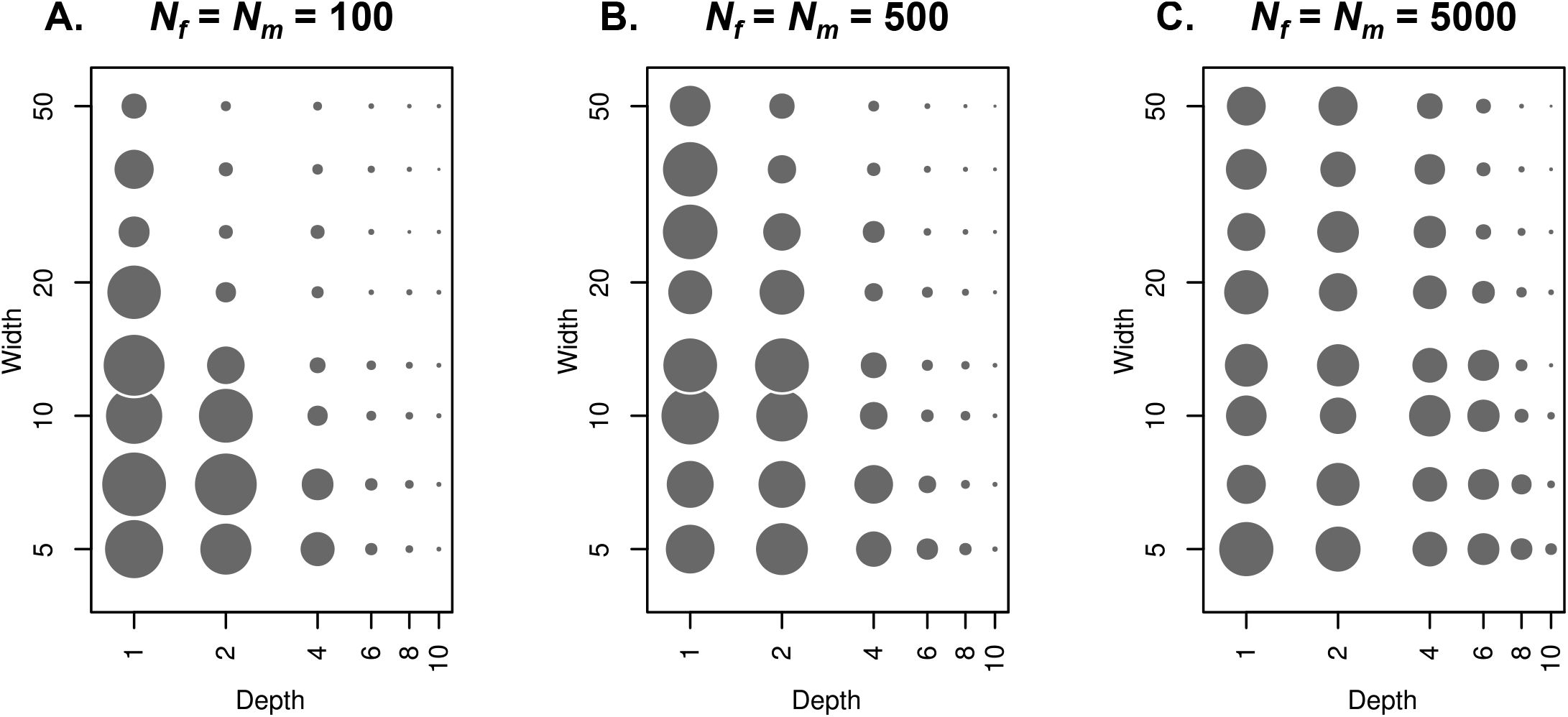
Power to infer population sizes as a function of pedigree width and depth. The surface of each dot represents the root mean squared errors (RMSE) over 10 simulations with population sizes (A) *N*_*f*_ = *N*_*m*_ = 100, (B) *N*_*f*_ = *N*_*m*_ = 500 and (C) *N*_*f*_ = *N*_*m*_ = 5,000. The RMSE is comprised between 1.681 × 10^−3^ for depth = 40 and width = 36 in (C) and 3.716 for depth = 1 and width = 7 in (A).

We next quantified the effect of pedigree depth and width (number of individuals at *g* = 0) on the accuracy of inferring population sizes. Maybe not surprisingly we found that much more information is contained in small but deep compared to large but shallow pedigrees (Figure 4). Indeed, increasing the width beyond just a handful of individuals seems to hardly reduce estimation accuracy except for very small population sizes, probably due to the oversampling effect described by Wakeley and Takahashi (2003).

The reason for this is that the number of individuals included in a completely resolved pedigree is growing rapidly with each generation going further back into the past (Derrida et al. 2000, Supplementary Figure S3), and so are the number of observed parent-offspring relationships informative about population size. Indeed, around 80% of the whole population is included in a complete pedigree of width 50 individuals at only few generations in the past, depending on the population size. At a depth of four, which we found to result in good estimates, about 7.5% or 750 individuals will be part of the complete pedigree of 50 individuals from a population of 10,000 individuals (Supplementary Figure S3).

## Discussion

Here we developed a model explicitly accounting for two sexes and overlapping generations. Under this model, genealogies follow a standard coalescent provided that time is rescaled appro-prietely and that expected coalesce times are much larger than the expected number of generations between parents and offspring, which is generally true under realistic parameter values. This new model allowed us to infer parameters jointly from genetic data and pedigree information. Using simulations we then show that including pedigree information not only improves the estimates of demographic parameters, but also allows to disentangle the effects of demographic and mutational processes on genetic diversity and hence to estimate these processes jointly. Indeed, our simulations show that with pedigree information of 50 individuals tracing back four generations is sufficient to obtain accurate joint estimates of male and the female effective population sizes, the proportion of overlapping generations and the mutation rate. Importantly, obtaining this amount of pedigree information is realistic for many populations of interest. For example, such pedigrees are available for several human populations (e.g., Hussin et al. 2015), for many domesticated animals breed of cattle and horses (e.g., Cunningham et al. 2001; Mc Parland *et al*. 2007) and for some wild animals (e.g., Clutton-Brock *et al*. 1982; Ellegren 1999).

However, we note that the amount of pedigree information required for accurate inference does depend on the population size with more data being required for larger populations. This stems from the fact that most of the information about the population size contained in a pedigree depends on the number of individuals sharing common ancestors. As a random sample is expected to contain less such individuals in a large population than in a smaller one, it will contain less information. Since the number of common ancestors increases more rapidly with the depth than the width of a pedigree, deep pedigrees of a few individuals contain much more information than shallow pedigrees of many individuals. As we discussed, the number of distinct ancestors in previous generations rapidly decreases with depth and reaches about 80% of the population within only few generations (Derrida et al. 2000, Supplementary Figure S3). However, these results consider a complete pedigree and are expected to be mitigated in presence of missing information.

Having only little pedigree data available will make it difficult to disentangle the effect of mutation and drift. A particular characteristic of such a situation is that the posterior distribution of the population sizes given the pedigree data alone will be very flat and often extend to very large population sizes. In such cases, the samples generated with our MCMC will likely not be distributed densely enough around the joint MLE to warrant accurate inference. In the extreme case of no pedigree information, the joint likelihood surface of the mutation rate and population sizes will form a ridge and the estimate produced by our stochastic inference method will single out a random combination not necessarily reflective of the true parameters. However, as we have show, already limited pedigree information of a few individuals over a few generations is sufficient to result in accurate inference.

While our theoretical and simulation results are very promising, we note that its application to real data may present some challenges. Firstly, the concept of generation, while convenient, is an artificial construct to discretize time that has little biological meaning for many long lived species. As a consequence, attributing the individuals of a pedigree to specific generations can be difficult. However, it is possible to extend our inference framework to also integrate over the attribution of individuals to generations as

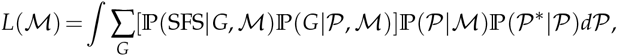

where we denote by 𝒫* the pedigree data without generation information (hence only relationships). Here, Pr(𝒫*|𝒫) = 1 if the pedigree 𝒫 is compatible with 𝒫*, that is, if all parents are from an older generation than all of their offspring and the most recently born individual is from generation 0, and Pr(𝒫*|𝒫) = 0 otherwise. Unfortunately, none of the parameters' MLE is trivial to derive because finding the maximum of this likelihood function implies finding the optimal set of pedigrees 𝒫. However, an MCMC method sampling such pedigrees can be envisioned to infer parameters under such an extended model.

Secondly, demographic events such as population size changes, migration between populations or complex mating systems, e.g., monogamy or harem models, may be needed to describe real populations. The introduction of such demographic events in the discrete generation pedigree model is fairly easy. For example, population size changes can be directly implemented in eq. 3 by using generation specific values of *N*_*f*_ and *N*_*m*_. The way demographic events shape coalescent processes is well described for many cases and they apply to our model if appropriately scaled. Complex mating systems or reproductive skew are well described for generation by generation models (Gasbarra *et al*. 2005) and some non-standard coalescent models are known to arise in these cases (Eldon and Wakeley 2006). In less extreme cases, specific mating systems can be well approximated by strongly skewed sex ratio (Nunney 1993) which our model already incorporates in its current form. It is important to note that inference under complex demographic scenarios is unlikely to work well with the MCMC-MLE approach introduced here as the pedigree may not have information for all demographic parameters, resulting in a bad proposal for the grid search. However, it is straight-forward to embed our model in an MCMC framework sampling from the joint posterior distribution.

In conclusion, we presented here a new model and some theoretical results on how to combine pedigree and genetic information for the inference of demographic and mutational process and showed that these processes can be disentangled if sufficient pedigree information is available. This is widely unexplored territory as most methods use individual or genealogy based models. But the availability of both pedigree and genetic data for many species, in particular domesticated animals, motivates the development of methods that combines such data. While an application to real data may pose additional challenges, our work is a first step towards such a method and extensions of our approach to more complex demographies and other features of real populations are readily possible. If done properly, the application of these to real data has the potential to give us deep insight into the mutational process in natural populations.

## Acknowledgements

We thank Dr. Christoph Leuenberger for helpful discussions on the early version of the model. FP and AT acknowledge support from the German Federal Ministry of Education and Research (BMBF) within the AgroClustEr ‘Synbreed-Synergistic plant and animal breeding’ (grant no. 0315528I). The work of S. M. Szilágyi was supported by the János Bolyai Fellowship Program of the Hungarian Academy of Sciences.

